# Electrical impedance spectroscopy with bacterial biofilms: neuronal-like behaviour

**DOI:** 10.1101/2023.11.24.568527

**Authors:** Emmanuel U. Akabuogu, Lin Zhang, Rok Krašovec, Ian S. Roberts, Thomas A. Waigh

## Abstract

Negative capacitance at low frequencies for neurons was first demonstrated in 1941 (Kenneth S. Cole) using extracellular electrodes. The phenomenon subsequently was explained by Cole using the Hodgkin-Huxley model and is due to the activity of voltage-gated potassium ion channels. We show that *E. coli* biofilms exhibit significant stable negative capacitances at low frequencies when they experience a small DC bias voltage in electrical impedance spectroscopy experiments. Using a frequency domain Hodgkin-Huxley model, we characterize the conditions for the emergence of this feature and demonstrate that the negative capacitance exists only in biofilms containing living cells. Furthermore, we established the importance of the voltage-gated potassium ion channel, Kch, using knock-down mutants. The experiments provide further evidence for voltage-gated ion channels in *E. coli* and a new, low-cost method to probe biofilm electrophysiology e.g. to understand the efficacy of antibiotics.

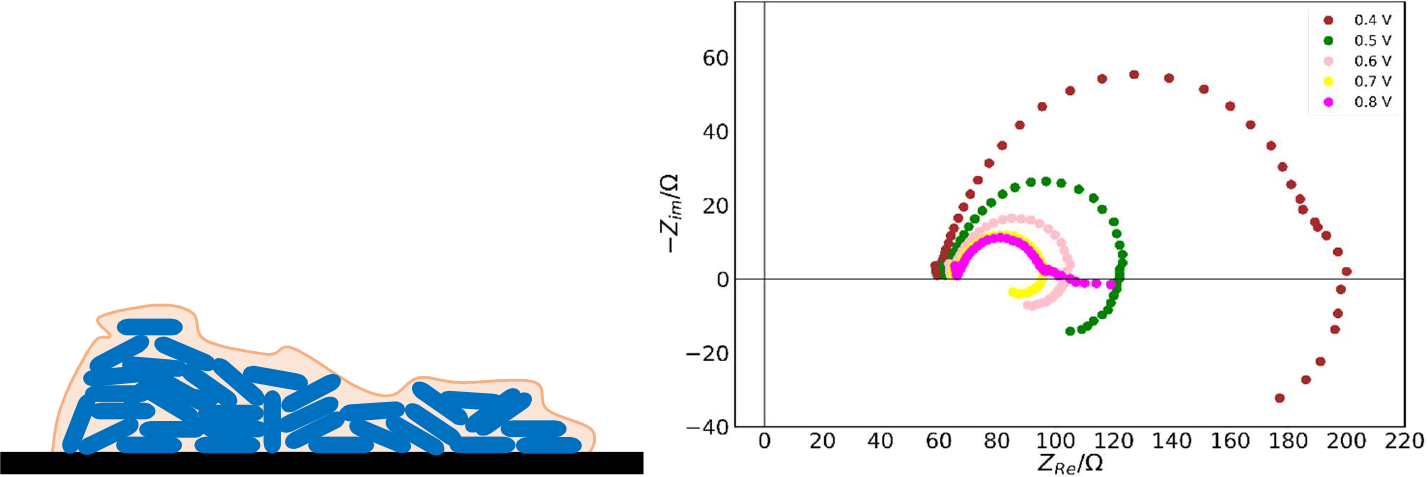

---

Transport of ions through voltage-gated ion channels in cellular membranes produces action potentials in neurons^1,2^. Historically, the electrical stimulation of neurons was investigated by Cole (1941), who was surprised to observe a large negative capacitance in electrical impedance spectroscopy experiments at low frequencies. The negative capacitance was eventually explained using calculations based on the Hodgkin-Huxley model^3,4^ and it is directly related to the activity of the voltage-gated potassium ion channels^3–6^.

Antibiotic-resistant bacteria currently present a huge problem to mankind^7,8^. Bacteria embedded in biofilms can be resistant to 2-3 orders of magnitude higher concentrations of antibiotics; a major factor contributing to the problem^9^. Electrical impedance spectroscopy has been previously applied to bacteria and bacterial biofilms^10–12^. However, the electrical impedance spectra looked similar to other organic layers and no specific neuronal-like behaviour has previously been observed.

Modern theoretical analysis shows the *negative capacitance* of neurons is crucial for their non-linear dynamics including bifurcations, stability and oscillatory spiking phenomena^13^. Negative capacitance also exists in inorganic systems, such as electrochemical cells^14^, memristors and solar cells^15–17^. For such systems, it can control catalytic processes, corrosion, memory effects and electrodeposition. In neurons, action potentials are caused by fluxes of Na^+^ and K^+^ ions through voltage-gated channels in the cell membranes^1,2^. Knowledge of this mechanism of neuronal dynamics has inspired synthetic devices which imitate action potentials in neurons. A specific example of such a device is the memristor, which has voltage and time-dependent resistance^18,19^. Memristors can adjust their resistance at a set voltage to mimick the action potential of neurons for neuromorphic computing. Memristors undergo resistive switching due to the amount of current that previously flowed, so they can act as memory devices in bioinspired networks^20,21^.

*Biofilms* are colonies of bacteria living together in extracellular polymeric substance (EPS) adhered to surfaces^9,22^. Bacteria in biofilms can have membrane potential dynamics similar to neurons^23–30^, although the hyperpolarization phenomena tend to be more varied than the stereotyped spiking events observed in neuronal cells. In the presence of external stimuli, such as metabolic stress^24^, photooxidative stress^25,30^, biosurfactants^31^ and external electrical stress^32^, cells in biofilms open their voltage-gated channels to allow the flow of cationic ions. In *B. subtilis* biofilms, the *YugO* channel controls the flux of potassium ions and is implicated in electrical signaling^24–26,28,33,34^. *E. coli* cells and biofilms engage in membrane potential spikes when stimulated by an electric field^32,35,36^. Our previous work also shows that the voltage-gated Kch channels in *E. coli* biofilms cause potassium ions to flow when exposed to light stress^23^. The flow of these ions triggers phases of hyperpolarization and repolarization which lead to electrical signaling across the biofilm. Neighboring cells in biofilms serve as conduits which propagate a wave of electrical signals via a fire-diffuse-fire mechanism.

In neuronal systems the effect of *negative capacitance* can be studied both in the time and frequency domains^37^. The time-domain version of the Hodgkin-Huxley (HH) model^6^ can be converted to the frequency-domain using Laplace transforms^3,5^. Equivalent circuits can be used to model the impedance spectra of samples under external voltage stimulation. The frequency-domain version of the HH model aids the study of ion channels and spike mechanisms, since it automatically averages over many thousands of stimulation events, providing succinct information not available with time-domain analysis and improved signal to noise. The frequency domain impedance response of the HH model for the squid axon under small amplitude perturbation was originally calculated by Cole^3,4^ and the analysis was recently extended by Bisquert et al^5^.

*Electrochemical impedance spectroscopy* (EIS) is a key experimental technique to probe the frequency domain electrical responses of electrochemical systems^38,39^. In EIS, small amplitude sinusoidal electrical perturbations are applied to thin layers of material attached to the electrodes and the frequency is varied over many decades. Analysis of the resulting complex impedance produces information on the electrochemical dynamics. The technique can also be used to decipher the dynamics of neurons to voltage stimulation^5^.

We characterized the impedance spectra of *E. coli* biofilms. We created an equivalent circuit (EC) model and subsequently established the frequency response of *E. coli* biofilms to small amplitude voltage perturbations.

Kch, a potassium ion channel in *E. coli* was discovered by Milkman in 1994 using comparative genetics techniques^40^. Its exact physiological role in *E. coli* is not completely understood, although there is circumstantial evidence that it is voltage-gated^41^ and previous evidence from our group shows that it is involved in electrosignaling in response to light stress^23^.

We used the method of P1 phage transduction to genetically move the Kch mutation from *E. coli* K-12 BW25113 Δ*kch*-mutant, a strain lacking the *kch* gene, into the wildtype (WT) DH5α. We confirmed the *E. coli* DH5α Δ*kch*-mutant phenotype using appropriate PCR primers, (Kch-F (5’-GTGAGTCACTGGGCTACATTCAAAC-3’) and Kch-R (5’-CTATTTTTGCGCCGATTCTTTAC-3’)). We grew the biofilms of the wildtype (WT) DH5α and DH5α Δ*kch* mutant strains on ITO electrodes. DH5α grows readily into a biofilm^42,43^ and removal of the Kch gene does not impede its growth^23,44,45^. Both strains exhibited similar growth curves determined using OD_600_ with a plate reader over 24 hr (Fig 1a). Next, we studied biofilm EIS after 24 hr growth, applying a 10 mV sinusoidal voltage over a frequency range of 2×10^-1^-10^5^ Hz with a 0 V DC voltage. DH5α exhibited distinct features in its Nyquist plot (Fig 1b) which were not observed in the DH5α Δ*kch* mutant (Fig 1c). Specifically, two distinct impedance arcs were observed in DH5α. The EIS spectra were stable and reproducible over the entire frequency range (Figs S1a and S1b). The impedance arc within the high-frequency region (Fig 1b, at the small impedances) was not seen in the DH5α Δ*kch* mutant (Fig 1c). We then analyzed the charge transfer resistance (*R*_*ct*_) within the biofilms. *R*_*ct*_ can be used to understand the rate of exchange of charge at the electrode/biofilm interface^46^. We designed an equivalent circuit (EC) model of the representative impedance spectra for both systems (Fig 1b inset and Fig 1c inset) and deduced the resistances *R*_*ctWT*_ and *R*_*ctMutant*_ for DH5α and the DH5α Δ*kch* mutant respectively. Remarkably, the charge transfer resistance of DH5α Δ*kch* mutant was *R*_*ctMutant*_ = 7500 ± 28 *Ω*, 15-fold higher than that for the wild type DH5α (Fig 1d), *R*_*ctWT*_ = 471 ± 4 *Ω*. This suggests the charge transfer between the electrode and the biofilm is much more efficient when Kch ion channels exist in the *E. coli* membranes.

**Fig 1:**
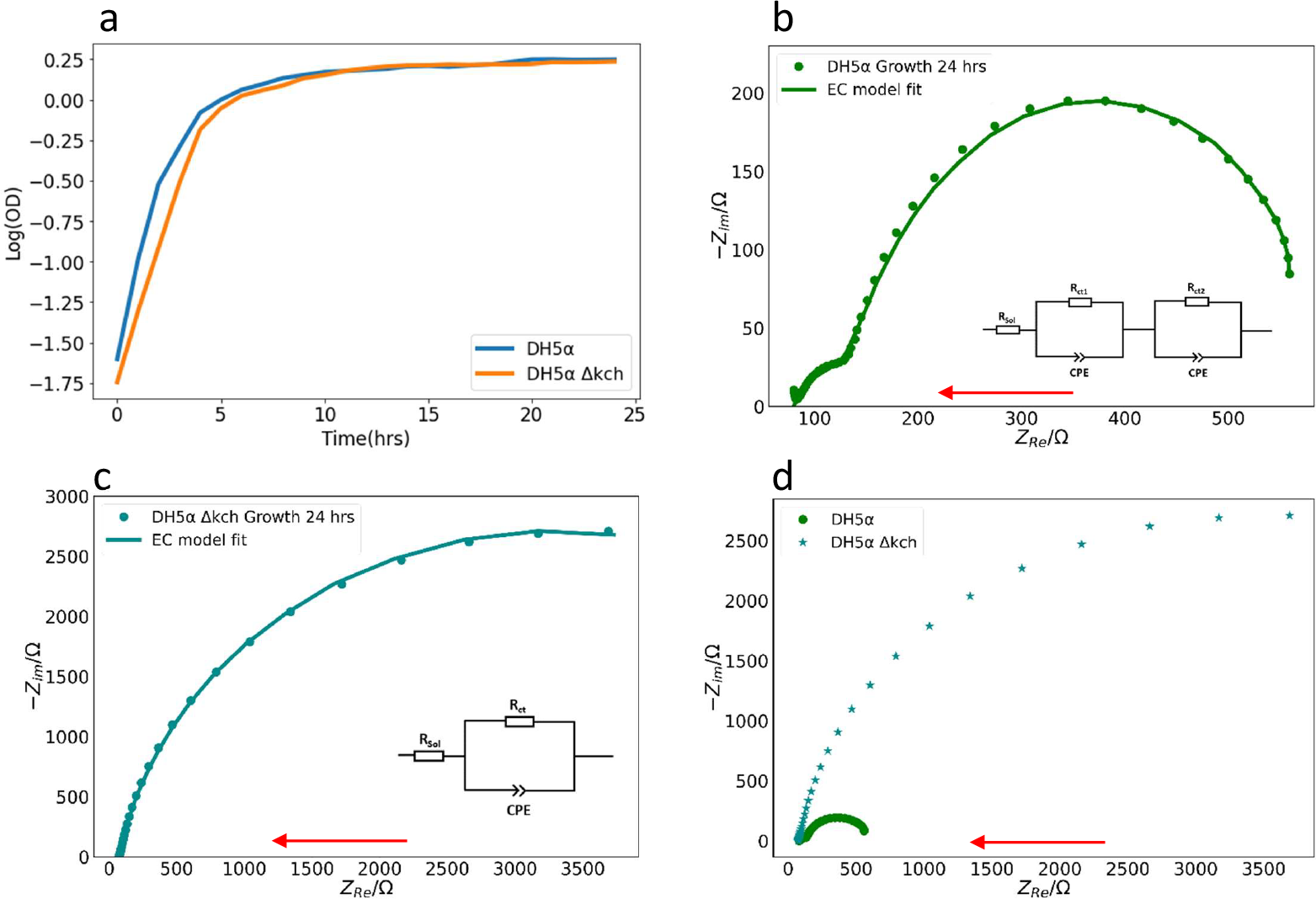
a) Growth curve for DH5α and DH5α Δ*kch* over 24 hrs from a plate reader. OD is the optical density. b) Representative impedance spectrum for DH5α showing the imaginary impedance (*–Z*_*im*_) plotted as a function of the real impedance (*Z*_*Re*_). Inset: Equivalent circuit model used to fit the data. The solid line represents the model fit using the equivalent electric circuit (EC) shown in the inset and the dots represent the experimental data. c) Representative impedance spectrum for DH5α Δ*kch*. Inset: Equivalent circuit model used to fit the data. The solid line represents the model fit using the EC shown in the inset and the dots represent the experimental data. d) Comparison between EIS spectra of both strains obtained after the 24 hr biofilm culture. All data were from at least three experimental replicates. Red arrows show the direction of increasing frequency.

We then monitored the growth-dependent change in EIS. We performed the EIS experiments for biofilms grown for 16-24 hr. The impedance plots for both strains revealed a steady decrease (Figs 2a, 2b) of the impedance arc i.e. larger more mature biofilms have lower impedances. The *R*_*ct*_ plot over time also confirmed the decreased charge transfer resistance of DH5α compared to DH5α Δ*kch* (Fig 2c).

**Fig 2:**
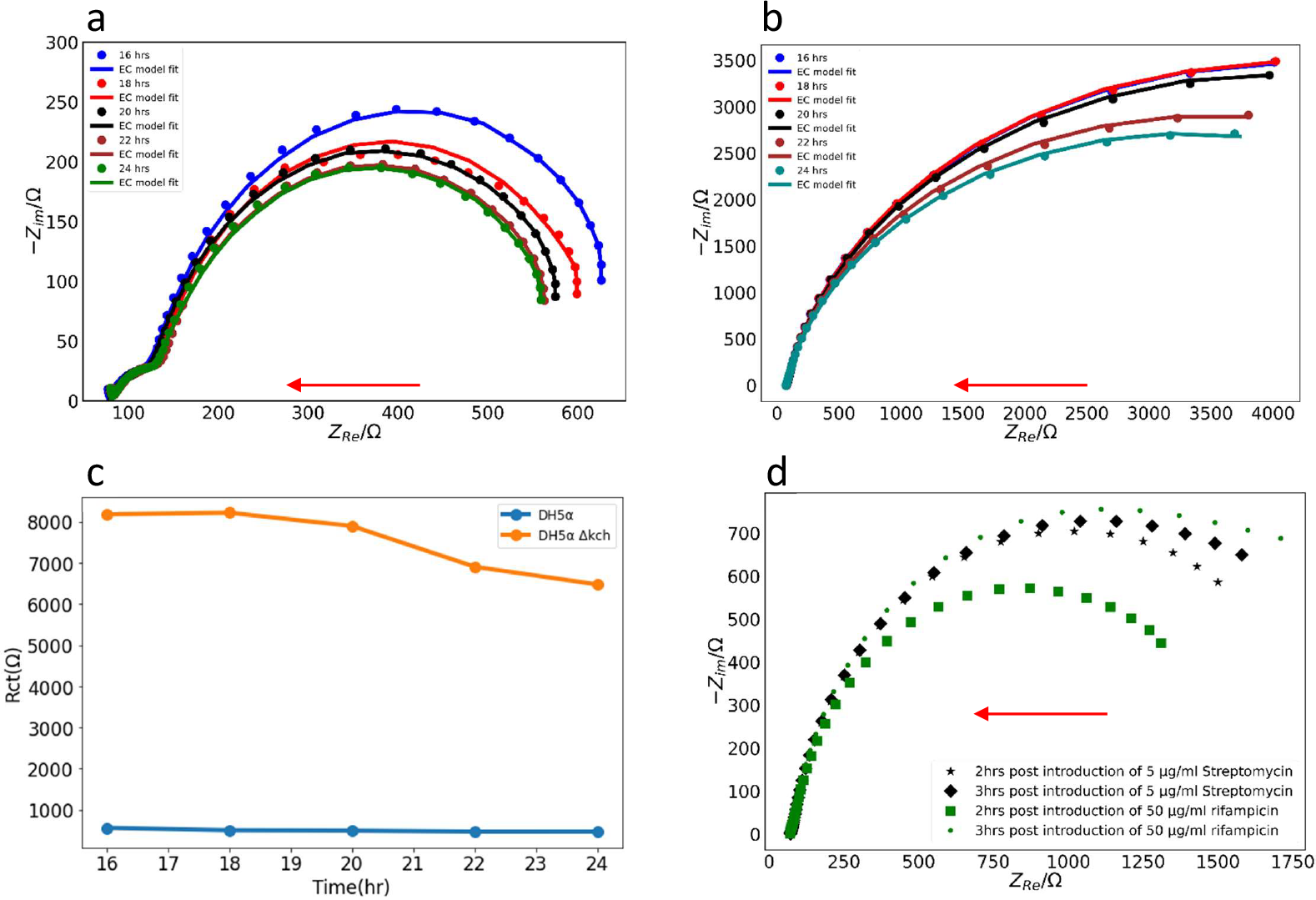
a) Variation of EIS spectra of DH5α biofilm as a function of growth time. Solid lines represent the model fits using the EC shown in the inset (Fig 1b) and dots represent the experimental data. b) Variation of EIS spectra of DH5α Δ*kch* biofilm as a function of growth time. Solid lines represent the model fits using the EC shown inset (Fig 1c) and the dots represent the experimental data. c) Time-dependent evolution of the charge transfer resistance in DH5α and DH5α Δ*kch* biofilm. For DH5α, the *R*_*ct2*_ from low impedance arc was used. d) Antibiotic-dependent changes in the EIS spectra obtained from DH5α biofilm. Red arrows show the direction of increasing frequency.

Next, we examined the effect of antibiotics on the observed impedance spectra of the DH5α strain. We employed two antibiotics, rifampicin and streptomycin at concentrations higher than their minimum inhibitory concentrations (SI). Rifampicin (net zero charge) inhibits bacterial RNA polymerase^47^, whereas streptomycin (positively charged) interferes with protein synthesis by ribosomes^48^. We observed an increase in the radius of the impedance arc and the *R*_*ct*_, as the concentration of the antibiotics increased i.e. the charge transfer was decreased by the antibiotics. Rifampicin showed a larger time-dependent increase in the *R*_*ct*_ compared to streptomycin (Fig 2d).

To explore the EIS spectra of *E. coli* biofilms in more detail, we conducted experiments at a range of DC bias voltages, 0.1-0.8 V. Experiments at different DC bias voltages help to monitor time-dependent memory effects in systems ^5,15,38,49^. When the *V*_*app*_ was increased from 0.1 V to 0.3 V, there was a corresponding steady decrease in the radius of the impedance arc (Fig 3a). The small semicircle observed at high-frequencies in the complex impedance plot at 0 V (Fig 1b and Fig 2a) disappeared. Furthermore, at a *V*_*app*_ of 0.4 V the low-frequency region becomes negative in the complex impedance plot i.e. a negative capacitance is observed. The impedance arc formed in the fourth quadrant of the impedance plot remained stable and persisted until the *V*_*app*_ approached 0.7 V. At a *V*_*app*_ of 0.8 V, the impedance arc in the fourth quadrant suddenly disappears, although the enlarged arc in the first quadrant remained (Figs 3a, 3b). A *V*_*app*_ of 0.4 V represents the threshold voltage for the appearance of negative capacitance. Further increases in the applied voltage *V*_*app*_ beyond 0.8 V yielded noisy data within the fourth quadrant.

**Fig 3:**
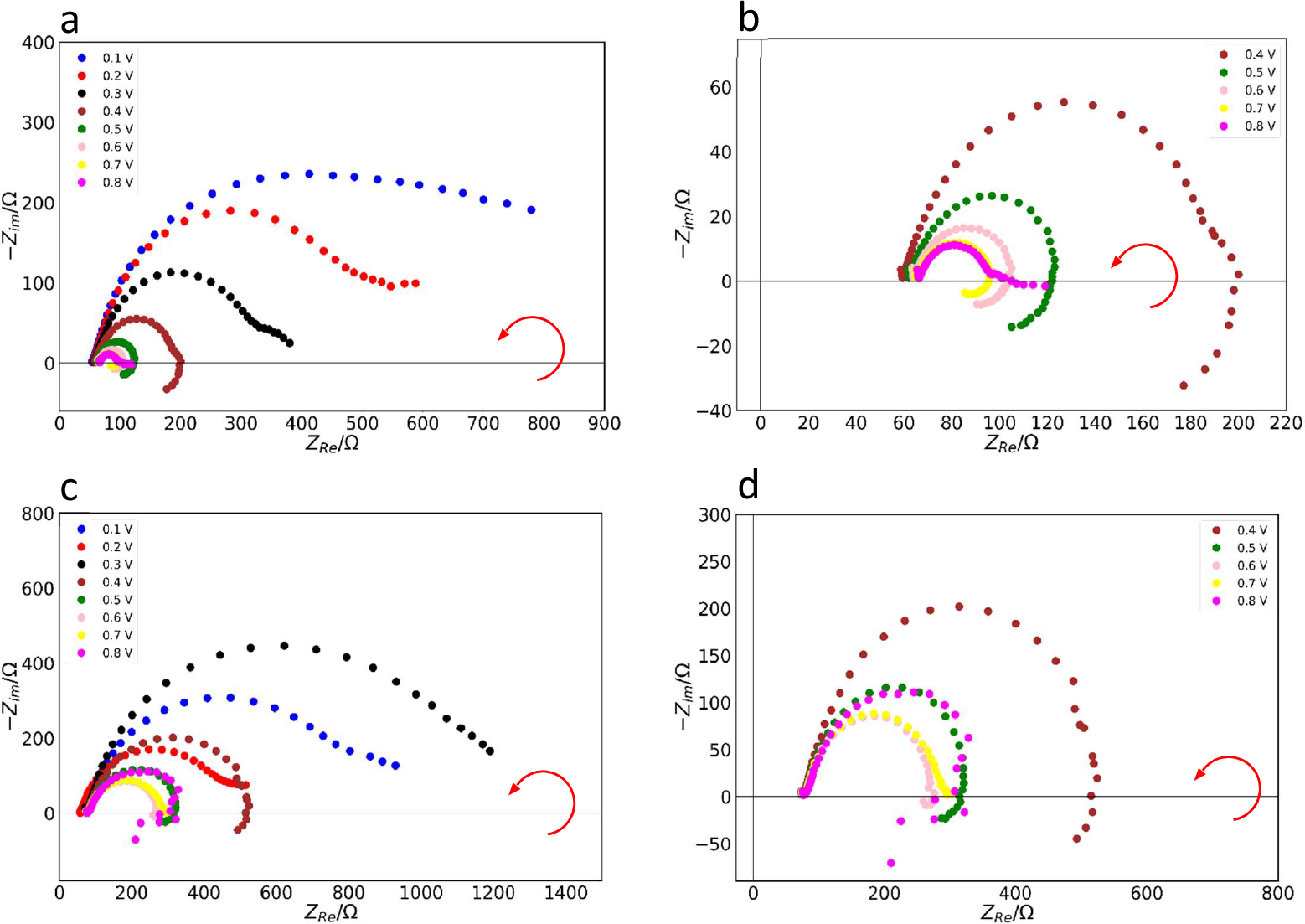
a) Electrical impedance spectra of DH5α biofilms at voltages, 0.1–0.8 V. b) Electrical impedance spectra of DH5α at DC bias voltages, 0.4–0.8 V, highlighting the negative capacitance. c) Electrical impedance spectra of DH5α Δ*kch* at DC bias voltages, 0.1–0.8 V. d) Electrical impedance spectra of DH5α Δ*kch* at DC bias voltages, 0.4–0.8 V, highlighting the negative capacitance. Red arrows show the direction of increasing frequency.

To understand the contribution of the Kch gene to the EIS data, we conducted EIS on the DH5α Δ*kch* biofilms under a variety of applied DC bias voltages. At a *V*_*app*_ of 0.1 V, we observed a similar-shaped spectra to the wildtype (Fig 3c). The impedance arc decreased in radius at a *V*_*app*_ of 0.2 V, but increased sharply at a *V*_*app*_ of 0.3 V. The impedance spectra transitioned to the fourth quadrant when the applied voltage was increased to 0.4 V i.e. a negative capacitance was observed. Compared with the wildtype biofilm, the radius of the impedance arc in the first quadrant of DH5α Δ*kch* was larger for each *V*_*app*_ (lower impedances, Fig S1c) and the impedance arc formed in the fourth quadrant disappeared at a *V*_*app*_ of 0.7 V (Fig 3d). The negative capacitance arc existed over a wider range of applied voltages in the wildtype than in the mutant (Fig 3 b and Fig 3d). The observed pattern of the EIS data for the DH5α and DH5α Δ*kch* biofilms at *V*_*app*_s of 0.4 – 0.7 V (Fig 3a-3d) and its stability, is consistent with the phenomenon of negative capacitance^3–5,37^. To create EIS data from DH5α biofilm which predominantly contains non-viable cells, the disinfectant Virkon was added (SI) and the negative capacitance behaviour was not observed (Fig S1d).

Motivated by work on the frequency domain response of the Hodgkin-Huxley (HH) model of the squid axon^4,5,50^ and the time domain version of the HH model for *E. coli* biofilms^23^, we developed equivalent circuits to characterize the frequency domain impedance response of the *E. coli* biofilms under small DC bias voltages (SI). Our focus was on the electrical impedance spectra produced at applied voltages of 0.4–0.7 V for the WT and 0.4–0.6 V for the Kch mutant where the negative capacitance phenomena were observed. Fig 4a is the predicted partial equivalent circuit for a single variety of voltage-gated ion channel^5^. The first branch contains the membrane capacitance (*C*_M_) and the second branch contains the resistance (*R*_*a*_) across *a*, a single variety of ion channel. The third branch is a parallel connection to a resistor-inductor (*R*_*x*_*L*_*x*_) circuit element which accounts for the inductive effect that produces the negative capacitance. *R*_*x*_ is the resistance across the gating variable *x* and *L*_*x*_ is the inductance across the gating variable *x*. The final branch consists of the resistance (*R*_*l*_) of the leak channel. While the partial circuit (Fig 4a) accounts for the membrane capacitance, ion conduction and the negative capacitance effects, more circuit elements are required to fit the experimental spectra (Fig 3 b and Fig 3d). Therefore, we created more complex minimal equivalent circuits to better describe the experimental data (Figs 4b, 4c). In our new equivalent circuit (Fig 4b), we added additional components to the single ion channel equivalent circuit (Fig 4a) to describe the contributions of the solution (*R*_sol_) and the interfacial surfaces (*R*_*ct*_) to the overall resistance. We also included a constant phase element (*CPE*) to account for any heterogeneity in the electrode surface^51^ and the formation of EPS in the biofilm over time^11^. This equivalent circuit (Fig 4b) described the impedance spectra of the voltage-gated ion channel(s) responsible for the negative capacitance observed in the DH5α Δ*kch* mutant (Fig 3d) and provides a description of voltage-gated channel(s) that are not Kch in the knockout mutant strain. We represent these voltage-gated ion channel(s) with the symbol *Q* and the cumulative resistance across those channels as *R*_*Q*_. *R*_*n*_ is the resistance associated with the gating variable *n* for the *Q* channel. The resistance *R*_*Q*_ and *R*_*n*_, and the time constant of the equivalent circuits were calculated as a function of the *V*_*app*_ (Figs 4d, Fig 4e, Table S1).

**Fig 4:**
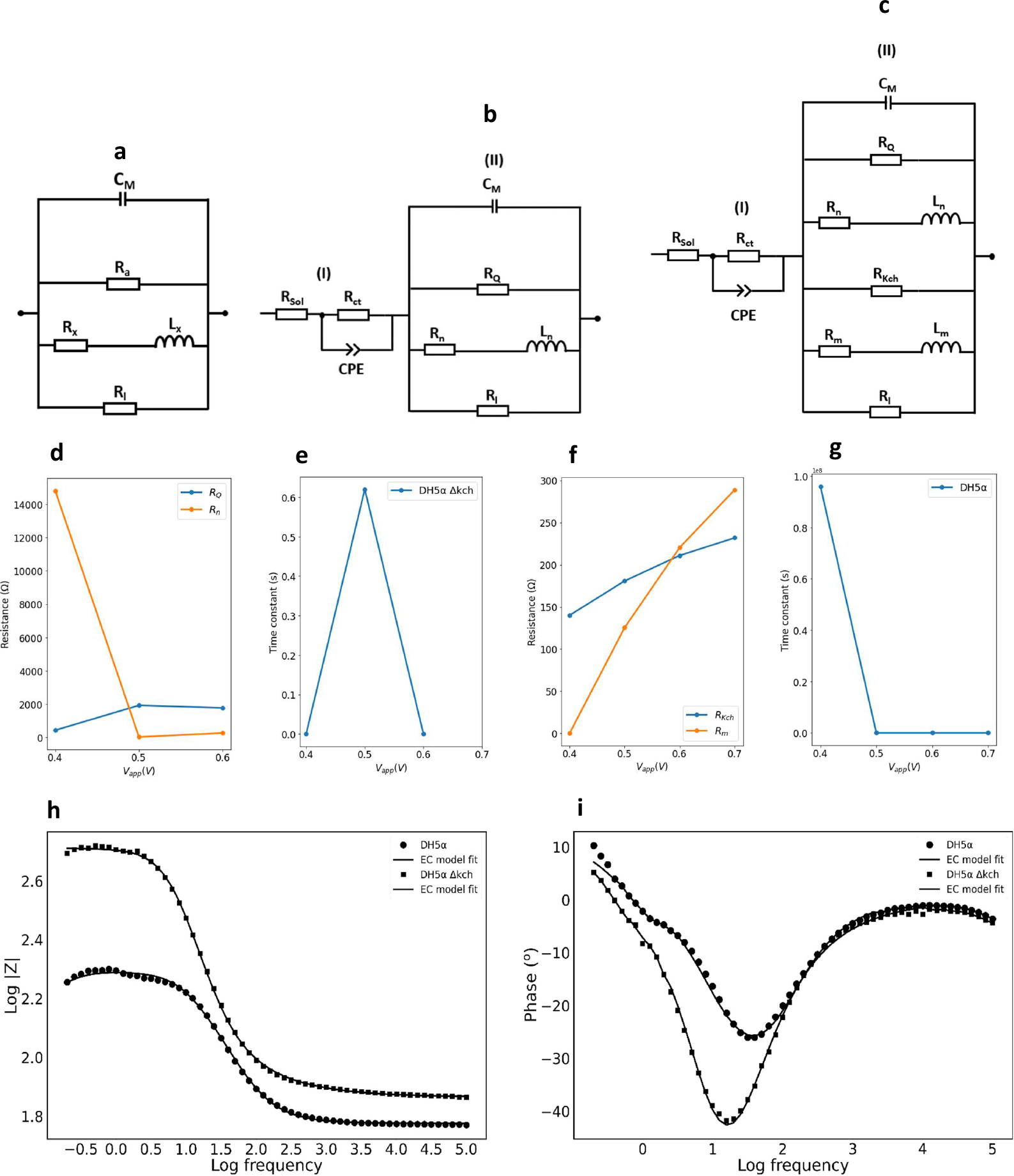
a) Partial equivalent circuit (EC) for the frequency domain Hodgkin-Huxley model for a single variety of voltage-gated ion channel^5^ . b) EC used to fit the EIS data for the *Q* channel in DH5α Δ*kch*, Fig 3d (0.4 – 0.6 V). c) EC used to fit the EIS data for the Kch channel in the wildtype, DH5α, Fig 3b (0.4 – 0.7 V). d) Values of the resistance for the range of applied voltages for the *Q* ion channel. Data obtained from Fig 3d using the EC fit of Fig 4b. R_Q_ and R_n_ are the resistances across the *Q* ion channel and its gating variable *n* respectively. e) Values of time constant for the *R*_*n*_*L*_*n*_ branch of the *Q* channel (Fig 4b) for the range of applied voltages. *L*_*n*_ is the inductance across the gating variable *n*. f) Values of the resistance for the range of applied voltages for the Kch ion channel of DH5α. Data obtained from Fig 3b using the EC fit of Fig 4c. *R*_*Kch*_ and *R*_*m*_ are the resistances across the Kch ion channel and its gating variable *m* respectively. g) Values of the time constant for the *R*_m_*L*_m_ branch of the Kch channel (Fig 4c, Fig 3b) for the range of applied voltages. *L*_*m*_ is the inductance across the gating variable *m*. h) Representative Bode plot (the change in the impedance modulus |*Z*| as a function of frequency) for both strains of *E. coli* at an applied DC bias of 0.4 V. Solid lines represent model fits using the EC (Fig 4c, DH5α and Fig 4b DH5α Δ*kch*) and the dots represent the experimental data. i) Representative Bode plot (the change in the phase as a function of frequency) for both strains at an applied DC bias of 0.4 V. Solid lines represent the model fits using the EC (Fig 4c, DH5α and Fig 4b DH5α Δ*kch*) and the dots represent the experimental data.

Next, we designed a complete equivalent circuit (EC) for the wildtype DH5α (Fig 4c). This EC incorporates the Kch channel and the other voltage-gated channels, *Q*. Data fits obtained for Kch ion channels in the WT, Fig 3b, using the circuit, Fig 4c, revealed increasing resistance values for *R*_*Kch*_ and the resistance of its gating variable, *R*_*m*_ (Fig 4f, Table S2). The time constant for the *RL* branch reveals a sharp drop as a function of applied voltage, followed by a plateau (Fig 4g). We confirmed that using the complete EC circuit (Fig 4c) to fit the DH5α Δ*kch* mutant spectra yields the same trend for *R*_Q_ and *R*_*n*_ (Fig S2a) as with using the EC of Fig 4b, i.e. the two models are mutually compatible.

We used Bode plots (Figs 3b, 3d) to resolve the dependence of the total impedance and phase on frequency. Fig 4h shows that for both strains, the impedance modulus had a slight rise within the low-frequency region and then a sharp drop within the mid-frequency region. However, across the entire frequency range, DH5α Δ*kch* exhibited a higher impedance modulus than the WT. Specifically, the impedance modulus obtained at the low frequency of 0.25 Hz is 3-fold higher than that for the WT at the same frequency. This difference in impedance modulus was consistent across the range of applied DC bias voltage (Table S3). This suggests that DH5α provide more efficient transport than DH5α Δ*kch* of cationic ions which is mediated by the potassium Kch ion channel at low frequencies. The negative capacitance effect was prominent within the low-frequency region (Figs 3b, 3d). This result agrees with the Nyquist plots (Figs 3a-3d) and the fit parameters from the model, in which the resistance across through the *Q* channel and its gating variable were high compared with those of the Kch channel (Tables S1, S2).

*E. coli* biofilms require higher values of the equivalent circuit elements to model their response to voltage stimulation than neurons, but biofilms contain many small bacterial cells whereas neuronal experiments are often performed on individual cells and the equivalent electrical components represent extensive physical variables. The resistance *R*_*Kch*_ across the potassium channel, Kch of *E. coli* biofilm varied between 140-232 ± 1 *Ω*, while the resistance *R*_*m*_ across its gating variable, *m*, was up to 289 ± 2 *Ω* (Fig 4f, Table S2). The biofilm inductance had values 0.94-53.8 ± 0.01 H (Table S3, Fig S2b). In biofilms, the time constant, *τ*_*m*_ = *L*_*m*_/*R*_*m*_ had values between (114 ± 3) x 10^-4^ and (9.6 ± 5.2) x 10^7^ across the range of applied voltages. In contrast, the predicted frequency domain EIS response of single neurons to voltage stimulation^5^ has typical resistance values, *R*_*K*_ and *R*_*n*_, ≤ 0.001 *Ω* and ≥ 0.05 *Ω* respectively, with the inductance not exceeding 2 H. Such high values of the inductance with neurons^4^ and biofilms are characteristic of their non-linear conductivities. The highest value for the time constant, *τ*_*n*_ = *L*_*n*_/*R*_*n*_ of the potassium channel of neurons was approximately 1 x 10^-3^ s ^5^ Neurons can therefore exhibit faster response times than *E. coli* biofilms by an order of magnitude.

Negative capacitance has been extensively linked to ion channel-mediated conductivity in different types of neuron^3–6,37^. The existence of the negative capacitance in both strains of bacterial biofilm in our study indicates the presence of voltage-gated ion channels in *E. coli* i.e. the Kch potassium channel and at least one other variety of ion channel. Our previous results show that biofilms of the DH5α Δ*kch* strain engage in ion-channel mediated membrane potential dynamics when under stress, although distinct changes were observed compared to the biofilm of the wildtype DH5α strain (the second hyperpolarization event in a novel two-spike phenomenon was lost in the Kch mutant)^23^. Our current study also demonstrates that the voltage-gated Kch potassium ion channel plays a major role in the electrical response of *E. coli* biofilms and the negative capacitance provides a unique signature of living *E. coli* in biofilms. Our results provide additional evidence of voltage-gating with the Kch potassium channel in *E. coli*. Negative resistance (as opposed to negative capacitance) is also observed in electrochemical impedance spectroscopy with neurons and it is attributed to the activity of sodium ion channels with both activation and inactivation gating^3,4^. No negative resistance was observed in the impedance spectra (2^nd^ and 3^rd^ quadrants) from bacterial biofilms indicating a lack of sodium-like ion channels.

The presence of the negative capacitance phenomenon in the biofilms of viable bacterial cells (Figs 3a-d) and its absence in the biofilm of non-viable cells (Fig S1d) could be used to quantify the viability of bacterial cells in biofilms. Portable EIS devices could be used with a small non-zero bias voltage to conduct LiveDead assays with bacterial biofilms. The method could be cheap, fast, high throughput and have a higher fidelity than previous methods. Specifically, EIS could prove to be less prone to artefacts than standard LiveDead assays using membrane-permeable/impermeable dyes^52–54^.

We characterized significant negative capacitance in bacterial biofilms at low frequencies i.e. a neuronal-like inductive effects in bacterial biofilms^4,5,37^. We believe this provides a unique insight into the electrophysiology of bacterial biofilms that should be explored with many more bacterial species e.g. *B. subtilis*^34^, *P. aeruginosa*^26^ and *S. lividans*^55 24,26,56^. Our electrical circuit models provided robust fits for the experimental data and enabled the extraction of key information about the physical processes in the biofilm. It has provided useful information on *E. coli* biofilm electrophysiology and crucially the technique does not require Nernstian voltage dyes that can be semi-quantitative and involve artefacts.

The human brain is very complex with a network 10^14^ synapses. Challenges therefore occur to build synthetic memristor networks which replicate synaptic responses of brains^57^. Synthetic biology techniques with bacterial biofilms could provide a basis for the design of simpler, cheaper, environmentally friendly and scalable bioinspired devices for neurocomputation.

## Acknowledgement

We thank Victor Martorelli for many useful discussions. R. K. is supported by the UKRI Future Leaders fellowship (MR/T021225/1). The authors would like to thank TETFund Nigeria and Abia State University Nigeria for E. A.’s PhD scholarship.

## Authors Contributions

E. A., L. Z., T. A. W. and I. R. conceptualized and designed the experiments. E. A. performed all experiments including strain generation. E. A. performed all data analysis. E. A. and T. A. W. wrote the manuscript with input from all authors. E. A. developed the models with input from L. Z. and T. A. W. E. A. performed the simulations.

