## Supplementary Information for "Electrical impedance spectroscopy with bacterial biofilms: neuronal-like behaviour"

### Bacterial biofilm culture

*E. coli* biofilm was used as the functional/active layer material. A day preceding the EIS experiment, the *E. coli* strains, DH5α and a DH5α Δ*kch* mutant were collected from a -80°C glycerol stock and streaked on an agar plate. On the day of the experiment, a single colony of the cell was transferred into a 10 ml Luria Broth (LB) media (Table S4). The glass universal containing the bacterial suspension was incubated overnight at 200 rpm at 37°C. The next day, 100 µl of the inoculum was pipetted into a fresh 10 ml LB and incubated in a shaking incubator at 200 rpm and at 37°C for 4.5 hours or $OD_{600} \approx 0.8$. Cells were subsequently adjusted to a starting concentration of 2 x $10^6$ CFU/ml for all the strains to ensure the same number of cells in all experiments. CFU/ml was calculated through the plate counting technique. Thereafter, 3 ml of the cell suspension was transferred onto an ITO electrode in the electrochemical cell. Cells were allowed 2 h to enhance cell attachment on the electrode surface at 37°C. Subsequently, 35 ml of fresh sterile LB was added to the cell suspension in the electrochemical cell to ensure proper electrical contact with the reference and counter electrodes. Cells were left to grow statically for 24 hrs at 37°C incubator. The optical density of the cells was measured using a spectrophotometer (JENWAY, Cole-Parmer UK). Each experiment was conducted in fresh electrolyte to eliminate the effect of culture media on the results.

### Working electrode cleaning, treatment, and functionalization

The transparent indium tin oxide (ITO) electrode (Sigma Aldrich) was employed as the conducting electrode. The ITO electrode easily integrates with bacterial substrates[1,2]. Prior to inoculation of the cell suspension on the ITO electrodes (2 x 2 $cm^2$), surface pretreatment was carried out to improve bacterial cell adhesion and biofilm cultivation[3,4]. The ITO surface was cleaned by sequentially sonicating for 10 min each in deionized water, acetone, and ethanol. The electrode was dried under nitrogen blow and subsequently treated with UV-ozone for 5 min. Following bacterial cell growth as described above, the biofilm cultivated for 24 hrs on the ITO serves as the electroactive layer material. The ITO electrode was not reused after each experiment.

### Device setup and electrochemical impedance spectroscopy

EIS experiments were conducted on a GAMRY potentiostat (GAMRY INSTRUMENTS, UK) using an electrochemical cell with a platinum counter electrode (CH Instruments Inc, US), an ITO working electrode and an Ag/AgCl reference electrode (CH Instruments Inc, US). After biofilm was grown for 24 h, EIS was performed at a small AC amplitude of 10 mV, a potential of 0 V and a frequency range from 0.2-$10^5$ Hz. Under the same conditions, EIS analysis was conducted at set times of 16, 18, 20, 22 and 24 h to monitor the spectral evolution during growth. The same procedure was employed for both *E. coli* DH5α and DH5α Δ*kch* mutant strains. To monitor the EIS of dead cells, the disinfectant Virkon (at the concentration of 1% (1:100)) was added after 24 hrs cell cultivation. The EIS data was collected 2 to 3 hrs after the addition of the disinfectant. This ensures no viable cells were left in the biofilm.

For the antibiotics experiments, we employed rifampicin at 50 µg/ml and streptomycin at 5 µg/ml. Data was collected 3 hrs after the addition of the antibiotics.

To study the effect of varying applied voltages, EIS experiments were performed at a constant small AC amplitude of 10 mV at different DC bias voltages ranging from 0.1 V to 0.8 V. The frequency sweep was maintained from 0.1 MHz to 0.2 Hz. For all experiments, Data acquisition, analysis and modeling were done with the GAMRY software (See Table 2).

### P1 phage transduction for the construction of *E. coli* DH5α Δ*kch* mutant strain

Using the PCR method, the Kch potassium gene was confirmed in a Keio collection WT strain, *E. coli* K-12 BW25113 strain[5] by amplification using the primers Kch-F (5'-GTGAGTCACTGGGCTACATTCAAAC-3') and Kch-R (5'-CTATTTTTGCGCCGATTCTTTAC-3'). Using the same primers, the Kch gene was not amplified in an *E. coli* K-12 BW25113 Δ*kch*-mutant strain. Subsequently, P1 lysate of the donor strain (*E. coli* K-12 BW25113 Δ*kch*-mutant) was prepared and transduced into the recipient (*E. coli* DH5α). Once more, the same primers were used to establish the absence of the Kch gene in our new strain, the *E. coli* DH5α Δ*kch* mutant.

### Model fit constants

**Table S1:**

| $V_{app}\ (V)$ | $R_Q$ (Ω) | $R_n$ (Ω) | $L_n$ (H) | $\tau_n = L_n/R_n$ (s) |
|---|---|---|---|---|
| $0.4$ | $429 \pm 3$ | $14800 \pm 100$ | $0.014 \pm 0.001$ | $(94 \pm 7) \times 10^{-8}$ |
| $0.5$ | $1920 \pm 20$ | $25 \pm 3$ | $16 \pm 2$ | $0.6 \pm 0.1$ |
| $0.6$ | $1770 \pm 10$ | $260.8 \pm 0.6$ | $0.00090 \pm 0.00005$ | $(34 \pm 2) \times 10^{-7}$ |

**Table S1:** The parameters extracted from model fits to the EIS spectra of *E. coli* DH5α Δ*kch* (Fig 3d) using the model in Fig 4b. $V_{app}$ is the applied DC bias voltage. $R_Q$ and $R_n$ are resistances across the $Q$ ion channel and its gating variable *n* respectively. $Q$ was used to represent all the voltage-gated ion channels present in the DH5α Δ*kch.* $L_n$ is the inductance across the gating variable *n.* $\tau_n$ is the time constant across the $RL$ branch of the circuit, Fig 4b.

**Table S2:**

| $V_{app}\ (V)$ | $R_{Kch}$ (Ω) | $R_m$ (Ω) | $L_m$ (H) | $\tau_m = L_m/R_m$(s) |
|---|---|---|---|---|
| $0.4$ | $140 \pm 1$ | $(6 \pm 3) \times 10^{-7}$ | $53.8 \pm 0.1$ | $(10 \pm 5) \times 10^{7}$ |
| $0.5$ | $181 \pm 1$ | $125.4 \pm 0.8$ | $0.94 \pm 0.01$ | $(750 \pm 9) \times 10^{-5}$ |
| $0.6$ | $211 \pm 3$ | $220 \pm 1$ | $2.68 \pm 0.02$ | $(121 \pm 1) \times 10^{-4}$ |
| $0.7$ | $232 \pm 1$ | $289 \pm 2$ | $3.2 \pm 0.1$ | $(111 \pm 3) \times 10^{-4}$ |

**Table S2:** Parameters extracted from the fits to the EIS spectra of *E. coli* DH5α (Fig 3b) using the model in Fig 4c. $V_{app}$ is the applied DC bias voltage. $R_{Kch}$ and $R_m$ are resistances across the $Kch$ ion channel and its gating variable *m* respectively. $L_m$ is the inductance across the gating variable *m.* $\tau_m$ is the time constant across the $RL$ branch of the circuit, Fig 4c.

**Table S3:**

| Strain | $\|Z\|_{freq=0.25\ Hz}$ at $V_{app} = 0.4\ V$ | $\|Z\|_{freq=0.25\ Hz}$ at $V_{app} = 0.5\ V$ | $\|Z\|_{freq=0.25\ Hz}$ at $V_{app} = 0.6\ V$ | $\|Z\|_{freq=0.25\ Hz}$ at $V_{app} = 0.7\ V$ |
|---|---|---|---|---|
| **DH5α Δ*kch*** | $508 \pm 4$ | $293 \pm 2$ | $258 \pm 1$ | – |
| **DH5α** | $188 \pm 2$ | $109.4 \pm 0.2$ | $92.4 \pm 0.1$ | $86.6 \pm 0.3$ |

**Table S3:** The impedance modulus at low frequency for both strains of *E. coli* biofilm for the range of applied DC bias voltages. Data was obtained at 0.25 Hz across the range of applied voltages with Bode plots similar to Fig 4h. The frequency of 0.25 Hz is equivalent to a log Frequency value of -0.6. $|Z|_{freq}$ represents the modulus of the impedance at a specific frequency. $V_{app}$ is the applied DC bias voltage.

**Table S4:**

| Media | Recipe | |
|---|---|---|
| **Luria Broth (LB)** | 10g/l NaCl, 5g/l yeast extract, 10g/l Tryptone. Distilled water to 1L | |
| **LB agar** | 10g/l NaCl, 5g/l yeast extract, 10g/l Tryptone, 15 g/l agar. | |
| **Experimental Models: Strains** | | |
| ***E. coli* DH5α** | Ian Robert's lab | |
| ***E. coli* DH5α (Δ*kch*)** | This study | |
| ***E. coli* BW25113 (JW1242-1)** | Keio Collection [8] | |
| ***E. coli* BW25113 (Δ*kch*)** | Keio Collection [8] | |
| **Software and Data Analysis** | | |
| **Python (Anaconda)** | Python | https://anaconda.org/ContinuumIO |
| **GAMRY** | GAMRY | https://www.gamry.com/support-2/software/ |
| **Electrochemical cell and Electrodes** | | |
| **Ag/AgCl** | CH Instruments Inc, US | |
| **Platinum (Pt)** | CH Instruments Inc, US | |
| **Transparent Indium Tin oxide (ITO)** | Sigma-Aldrich | |
| **Electrochemical cell** | GaossUnion, China | |
| **GAMRY Potentiostat** | GAMRY Instruments Inc, US | |

**Table S3:** Media recipe, bacterial strains, software and EIS components used in experiments on *E. coli* biofilms.

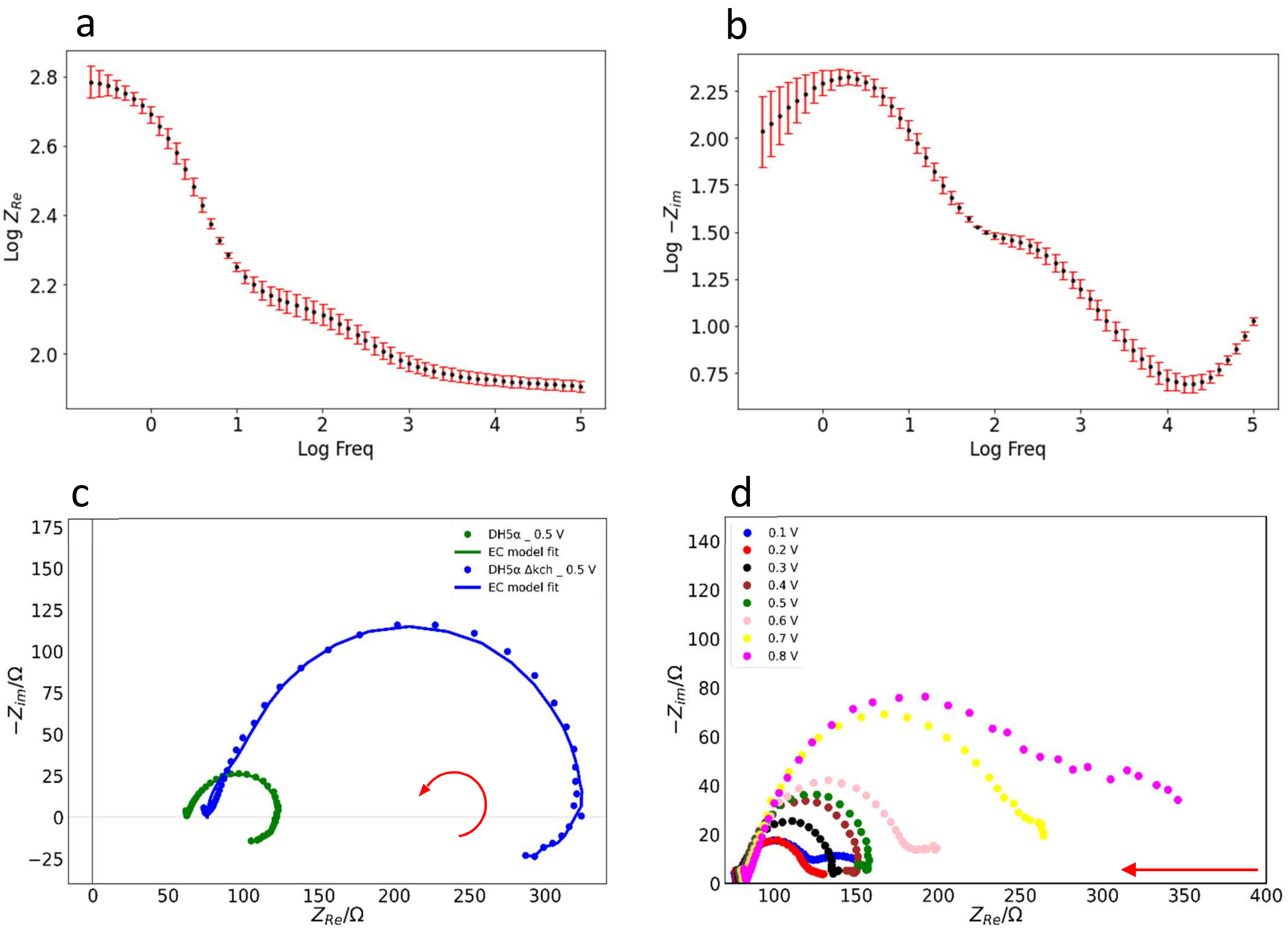


**Fig S1:** (a) The real impedance ($Z_{Re}$) plotted as a function of frequency from experiments with the wildtype DH5α biofilms (Fig 1b) (mean ± SD for three repeats). b) Imaginary impedance ($-Z_{Im}$) as a function of frequency from the wildtype DH5α biofilms (Fig 1b). c) Representative data showing that our proposed minimal equivalent circuit provides a good fit to experimental data with the imaginary ($-Z_{Im}$) plotted as a function of the real impedance ($Z_{Re}$). The equivalent circuits of Figures 4b and 4c were used for the mutant and wildtype respectively. The solid lines are model fits and the dots are the experimental data at the applied DC bias voltage of 0.5 V. The plot also compares differences in the impedance arcs at similar DC bias voltages for both strains. Each point shows a separate frequency. d) EIS data for DH5α biofilm with no viable cells (subjected to Virkon for 2 hours) for different DC bias voltages showing the complex impedance ($-Z_{Im}$) as a function of the real impedance ($Z_{Re}$). Each point shows a separate frequency. Red arrows show the direction of increasing frequency.

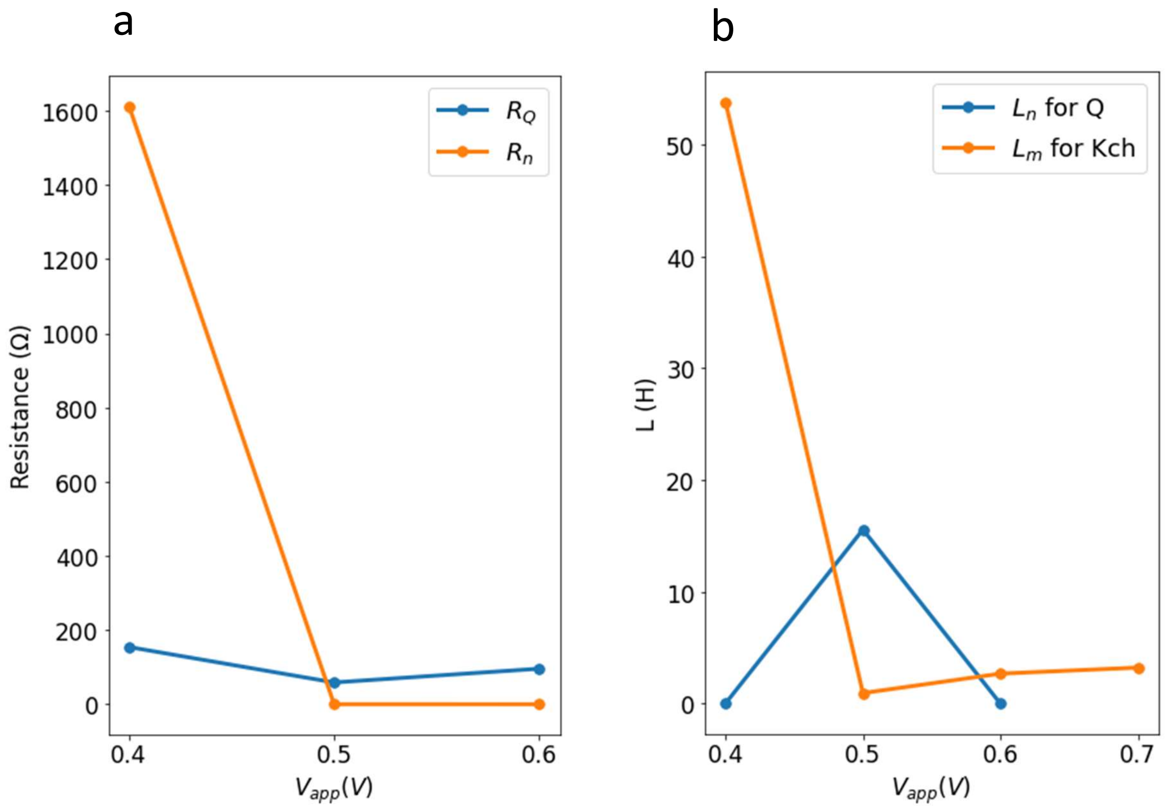


**Fig S2:** a) Values of the resistance for the Kch mutant as a function of the DC bias voltage $(V_{app})$ from the EIS data shown in Fig 3d obtained using the complete equivalent circuit model shown in Fig 4c. $R_Q$ and $R_n$ are resistances across the $Q$ ion channel and its gating variable *n* respectively. b) Values of the inductances $(L)$ as a function of the DC bias voltage $(V_{app})$ for both *E. coli* biofilm strains using their respective equivalent circuits models. $L_n$ and $L_m$ are the inductances across the gating variables *n* and *m* of the $Q$ and $Kch$ ion channels respectively.

### Mathematical model of the frequency domain impedance response of *E. coli* biofilms

A single channel time-dependent Hodgkin-Huxley model[6] was developed to explain the stress-dependent electrical signalling in *B. subtilis* biofilms[7]. We extended this model to understand the two-channel mediated membrane potential dynamics in *E. coli* biofilms in response to blue light stress[8]. In the current work, we used the same two-channel Hodgkin-Huxley model as our previous study in *E. coli*[8] to understand electrical impedance spectroscopy measurements with *E. coli* biofilms.

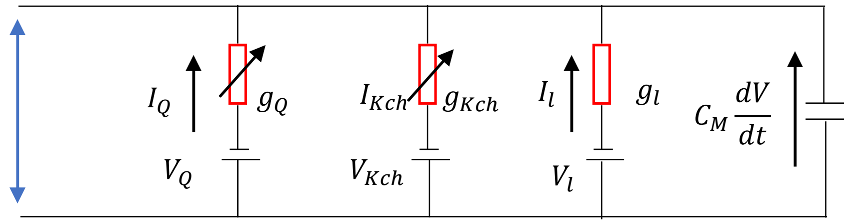


**Fig S3:** A Hodgkin-Huxley equivalent circuit model for the time-dependent conductance of *E. coli* biofilms[8]. $g_Q$, $g_{Kch}$ and $g_l$ represent the conductance of the, $Q$, $Kch$ and the leak channel $l$ respectively. $V_q$ , $V_{Kch}$ and $V_l$ represents the Nernst potentials for the $Q$, $Kch$ and leak ions respectively. $I_Q$, $I_{Kch}$ and $I_l$ are currents through the $Q$, $Kch$ and leak channels respectively. $C_M(dV/dt)$ represents the capacitive current across the membrane.

Fig S3 shows the equivalent circuit used in the HH model of *E. coli* biofilms in which the bacteria are modelled with two ion channels Kch and Q. The currents through the ionic channels (the equivalent resistors) and the cell membrane (the equivalent capacitor) in Fig S3 are

$$I_c = C_M \frac{dV_M}{dt} , \quad (1)$$

$$I_Q = g_Q(V_M - V_Q) , \quad (2)$$

$$I_{Kch} = g_{Kch}(V_M - V_{Kch}), \quad (3)$$

and

$$I_l = g_l(V_M - V_l). \quad (4)$$

where $g_j = 1/R_j$ is the conductance of the $j$ ion channel. Thus $g_Q$, $g_{Kch}$ and $g_l$ are the conductances of the, $Q$, $Kch$ and leak channel $l$ respectively. $V_q$ , $V_{Kch}$ and $V_l$ represent the Nernst potentials for the $Q$, $Kch$ and leak ions respectively. $Q$ represents all voltage-gated channels other than the $Kch$ ion channel which are in the DH5α Δ*kch* mutant. $V_M$ is the resting Nernst potential.

We assumed that both positively charged ion channels, $Q$ and $Kch$, have four subunits which are their four activation gates. The variable *n* represents the gating variable for the $Q$ channels and *m* is the gating variable for $Kch$ channel. We can therefore rewrite eqns 2, 3 and 4 as

$$I_Q = \frac{1}{R_Q} n^4 (V_M - V_Q) \, , \qquad 5$$

$$I_{Kch} = \frac{1}{R_{Kch}} m^4 (V_M - V_{Kch}) \, , \qquad 6$$

$$I_l = \frac{1}{R_l} (V_M - V_l) \, . \qquad 7$$

The fraction of time each channel is open can then be represented by

$$\frac{dn}{dt} = \alpha_n(S)(1-n) - \beta_n \; , \qquad 8$$

and

$$\frac{dm}{dt} = \alpha_m(S)(1-m) - \beta_m \; , \qquad 9$$

for the $Q$ and $Kch$ channels respectively. $S$ represents the stress level[8–10], e.g. due to voltage stimulation or light stress. $\beta_n$ and $\beta_m$ stand for the rate at which the open channels close and they are stress-dependent. $\alpha_n$ and $\alpha_m$ are the rates at which closed channels open and they are stress-dependent. $R_Q$, $R_{Kc}$ , and $R_l$ are resistances across the $Q$, $Kch$ and $l$ respectively.

Busquert et al. provided a small perturbation AC model for the frequency domain impedance response of the standard HH model for a neuron (with both potassium and sodium ion channels) which applies to each branch of the neuronal equivalent circuit[11]. Based on our experimental observations, we therefore extended this version of the HH model to develop a minimal model for the frequency domain impedance response of *E. coli* biofilms under small AC perturbations i.e. we include two potassium-type ion channels because negative capacitances are observed, but no negative resistances.

To model AC perturbations of the HH model, a Laplace transform of eqns 1, 5, 6 and 7 yields

$$\tilde{I}_C = sC_M\tilde{V}_M \,, \qquad 10$$

$$\tilde{I}_Q = \frac{1}{R_Q}4\bar{n}^3\left(\bar{V}_M - V_Q\right)\tilde{n} + \frac{1}{R_Q}\bar{n}^4\tilde{V}_M \,, \qquad 11$$

$$\tilde{I}_{Kch} = \frac{1}{R_{Kch}}4\bar{m}^3(\bar{V}_M - V_{Kch})\tilde{m} + \frac{1}{R_{Kch}}\bar{m}^4\tilde{V}_M \,, \qquad 12$$

$$\tilde{I}_l = \frac{1}{R_l}\tilde{V}_M, \qquad 13$$

where $\tilde{I}_C$ , $\tilde{I}_Q$ and $\tilde{I}_{Kch}$ are the current across the membrane, the $Q$, $Kch$ and $l$ respectively. $\tilde{V}_M$ is the resting Nernst potential. The Also $\tilde{n}$, $\tilde{m}$ are the perturbed gating variables. The tilde sign (~) represents the small perturbation value. $\bar{n}$ and $\bar{m}$ represent the value of the gating variables at steady state. $\bar{V}_M$ is the value of the Nernst potential at a steady state. The overbar (¯) represents the value at a steady state.

The Laplace transform of the gating variables *n* and *m* which are defined eqns 8, 9 and which also appear in eqns 11, 12 yields

$$s\tilde{n} = \left[\frac{\partial\bar{\alpha}_n}{\partial V_M}(1-\bar{n}) - \frac{\partial\bar{\beta}_n}{\partial V_M}\bar{n}\right]\tilde{V}_M - \left(\bar{\alpha}_n + \bar{\beta}_n\right)\tilde{n} \qquad 14$$

$$s\tilde{m} = \left[\frac{\partial\bar{\alpha}_m}{\partial V_M}(1-\bar{m}) - \frac{\partial\bar{\beta}_m}{\partial V_M}\bar{m}\right]\tilde{V}_M - \left(\bar{\alpha}_m + \bar{\beta}_m\right)\tilde{m} \qquad 15$$

The impedance Z for the frequency domain response is

$$Z = \frac{\tilde{V}_M}{\tilde{I}_M} = \frac{\tilde{V}_M}{\tilde{I}_C + \tilde{I}_Q + \tilde{I}_{Kch} + \tilde{I}_l}. \qquad 16$$

Substituting the eqns 10-15 into 16, we obtain

$$Z = \left[ sC_M + \frac{1}{R_Q} + \frac{1}{R_n + sL_n} + \frac{1}{R_{Kch}} + \frac{1}{R_m + sL_m} \right]^{-1} \quad 17$$

where $s$ is the Laplace frequency, $\omega$ is the frequency and $s = i\omega$. The overbar denotes the values at a steady state while the tilde represents the small perturbation. $R_n$ and $R_m$ denote the resistances through the gating variable *n* and *m* respectively. $L_n$ and $L_m$ are the inductance across the gating variable *n* and *m* respectively.

Eqn 17 allows the frequency domain response of the electrical equivalent circuit Fig 4c of the *E. coli* biofilm to be calculated.

Each circuit element which depends on the voltage can then be deduced as stated below:

$$R_Q(\bar{V}_M) = \frac{R_Q}{\bar{n}^4} \quad 18$$

$$R_n(\bar{V}_M) = \frac{R_Q}{4\bar{n}^3(\bar{V}_M - V_Q)\tau_n \left[ \frac{\partial \bar{\alpha}_n}{\partial V_M}(1 - n) - \frac{\partial \bar{\beta}_n}{\partial V_M}\bar{n} \right]} \quad 19$$

$$L_n(\bar{V}_M) = R_n \tau_n \quad 18$$

$$R_{Kch}(\bar{V}_M) = \frac{R_{Kch}}{\bar{m}^4} \quad 20$$

$$R_m(\bar{V}_M) = \frac{R_{Kch}}{4\bar{m}^3(\bar{V}_M - V_{Kch})\tau_m \left[ \frac{\partial \bar{\alpha}_m}{\partial V_M}(1 - m) - \frac{\partial \bar{\beta}_m}{\partial V_M}\bar{m} \right]} \quad 21$$

$$L_m(\bar{V}_M) = R_m \tau_m \quad 22$$

where $\tau_n$ and $\tau_m$ are the relaxation time constants for the gate variables, *n* and *m* respectively. $\bar{\alpha}_n$ and $\bar{\beta}_n$ are the values of the gating variable *n* at steady state. $\bar{\alpha}_m$ and $\bar{\beta}_m$ are the values of the gating variable *m* at steady state.
